# Extracellular vesicle packaging of a supercluster-encoded antimicrobial combination with enhanced potency against fungal biofilms

**DOI:** 10.64898/2026.10.25.761712

**Authors:** Kirsten J. Meyer, Valerie Chow, Elaina Piliouras, Yunjin Lee, Hanna Ostapska, Oliver Ojeda Valencia, Ines Ben Makhlouf, Nicole Robbins, Leah E. Cowen, Justin R. Nodwell

**Author notes:** KJM – Department of Biology, University of Waterloo, HO – Department of Microbiology, University of Montreal, IBM – Division of Chemistry and Chemical Engineering, California Institute of Technology.

## Abstract

Bacterial specialized metabolites are bioactive compounds that have been mined extensively as antibacterials, antifungals, and other drugs. They are widely used as experimental probes of complex biological systems. Since their discovery, their functional characteristics have typically been investigated as single, soluble compounds. The discoveries in this work expand this model. First, we report the co-packaging in extracellular vesicles (EVs) of two specialized metabolite complexes, candicidin and antimycin A, the products of a supercluster of biosynthetic genes, by a *Streptomyces* soil isolate. Secondly, we report that the products of this supercluster have synergistic antimicrobial activity against fungal biofilms, whereas biofilms are tolerant to the individual compounds. Finally, we show that co-packaging these compounds in EVs ensures simultaneous delivery with enhanced potency. This research advances understanding of the elegant complexity that shapes the evolution and function of bacterial specialized metabolites. In particular, the enhanced antibiofilm activity of co-packaged metabolites compared to individual soluble compounds is notable given the relevance of biofilms to microbial ecology and to pathogenesis. Our work supports the evolution of coordinated activity of specialized metabolites to counter conditional environmental challenges, and highlights co-packaging in EVs as one mechanism for secretion and co-delivery of combinations of bioactive molecules.

## INTRODUCTION

Many bacteria secrete synthetically complex specialized metabolites that have high affinities for specific biological targets including proteins, nucleic acids, lipids, and carbohydrates^1–5^. These interactions confer potent bioactivities, often with implications for communication and competition with neighboring cells. Isolation of these specialized metabolites continues to provide exquisitely selective chemical probes and lead molecules for research and development, as well as the majority of novel antimicrobials^6–10^. Given the enormous breadth of microbial and specialized metabolite diversity, our understanding of the complex regulation, secretion, and transport mechanisms of these metabolites by microbes is still limited^11–17^.

The genes responsible for the production of specialized metabolites are organized into biosynthetic gene clusters (BGCs)^3^. Genomic analysis has consistently found that actinomycete bacteria, in particular the genus *Streptomyces*, have the greatest diversity and number of BGCs^1^. There are over 7000 unique gene cluster families across the genera, with each species of *Streptomyces* harbouring 20-30 BGCs^1,15^. There is strong evidence for coordinated regulation of multiple gene clusters, although our understanding of this is limited to a few species^11,18–22^. Contingency and synergy have been proposed as driving forces for the evolution of coordinated regulation^18^. “Contingency”, suggests that producing multiple metabolites with different bioactivities covers more than one possible environmental condition. “Synergy”, suggests that bioactivity can be enhanced by combining compounds with different targets and/or mechanisms. An increasingly observed mechanism for coordinated regulation and activity of two or more metabolites is the genomic BGC “supercluster”, where the BGCs for chemically-distinct metabolites are co-localized^18–20,23,24^. These supercluster metabolites are of great interest as several have shown enhanced antimicrobial activity particularly in the context of bypassing or preventing resistance mechanisms in target microbes^18–20^.

The first supercluster identified, and to date the most impactful, is the β-lactamase inhibitor clavulanic acid co-produced with cephamycin C^25,26^. Given the widespread production of β-lactamases in environmental bacteria^27^, it seems likely that co-production of clavulanic acid with cephamycin C extends the bioactivity of cephamycin C for the producing *Streptomyces*^26^. Clavulanic acid was clinically developed in combination with amoxicillin in the 1980s, setting the benchmark for several decades of successful β-lactamase inhibitor development in the fight against antibiotic resistance^19,28^. A recent example adding to the small but expanding collection of characterized superclusters is the identification of polycyclic tetramate macrolactam (PoTeMs) metabolite genes co-localizing with a variety of BGCs in diverse *Streptomyces sp.* These PoTeMs can permeabilize membranes, countering the intrinsic resistance of the cell envelope as a barrier to entry for their partner metabolites^24^. A final example is the interesting report of a widespread megacluster consisting of a streptavidin-flanked BGC encoding four metabolites that each target different steps in biotin metabolism^23^. The encoded metabolites exhibit profound synergy against *Mycobacteria* and Gram-negative bacteria in physiologically and ecologically relevant biotin-limited conditions^23^. This megacluster highlights the interplay between environmental conditions and the contingent benefit specialized metabolites may provide to the producing organisms^18,29^.

A major challenge for ensuring the biological utility of such combinations is ensuring that the partner compounds are produced, secreted, and delivered to their target cells simultaneously. This is particularly challenging because different compounds will have distinct diffusion characteristics. Indeed, the typical model of specialized metabolite release is by direct secretion across the cell membrane by efflux pumps and transporters and into the extracellular space as free chemicals^3^. Diversifying this model, we and others have found that many *Streptomyces* species can package antimicrobials into extracellular vesicles (EVs)^30–34^. EVs are 20-200 nm lipid bound entities that have emerged as mediators of inter-cellular interactions, within and between species and even kingdoms^35,36^. Bacterial EVs are formed from the outer membrane in Gram-negative^37,38^, and from the cellular membrane in Gram-positive bacteria particularly at areas of cell-wall turnover^39,40^. Given the evidence that EVs can serve to protect their molecular cargo, alter diffusion, and play a key role in delivery^41–45^, we wondered whether EVs can serve as a delivery mechanism for specialized metabolite combinations.

In this work, we report the packaging of a combination of candicidin and antimycin A, derived from a conserved genomic supercluster^46^, into extracellular vesicles. This occurs in a soil isolate, WAC276, part of the *Streptomyces albidoffavus* clade. Candicidin is a well known polyene antifungal, originally described by Waksman in 1952^47,48^. Antimycin A potently inhibits eukaryotic cytochrome bc1, complex III of the electron transport chain^49^. Candicidin was recently identified in EVs from *S. albidoffavus* S4^34^, but not co-packaged with antimycin A. Importantly we find both candicidin and antimycin A co-purifying in EVs, and we show that this combination in the EVs exhibits contingent synergy against fungal biofilms of *Cryptococcus neoformans* and *Candida albicans*. *C. neoformans* is an environmental yeast and opportunistic human pathogen^50^, and *C. albicans* is the leading cause of fungal infections in humans^51^. Microbial biofilms are attached multicellular communities surrounded by an extracellular matrix of biopolymers. Biofilms are notoriously tolerant to individual antimicrobials and a widespread problem in the clinic^52^, but also a common feature of microbial habitats in the environment^53^. We demonstrate that vesicle packaging of candicidin and antimycin A ensures co-delivery of these compounds that otherwise have markedly different diffusion properties. The EVs are able to traverse the biofilm and achieve membrane fusion with the fungal hyphae. Our results support contingency and synergy as selective pressures for the evolution and function of specialized metabolite combinations in nature. They further demonstrate how these privileged natural combinations can counter tolerance mechanisms in microbes. Perhaps most importantly, we show that packaging in EVs supports combinatorial antimicrobial activity in environmentally and clinically relevant contexts such as a fungal biofilm.

## RESULTS

### WAC276 vesicles contain candicidins and antimycins, products of a biosynthetic supercluster

To identify actinomycete EVs packaging cocktails of specialized metabolites, we fractionated EV cargo using reverse-phase chromatography and tested for distinct zones of antimicrobial activity. In previous work, we showed that the actinomycete strain WAC276 is a producer of potent antifungal EVs (Fig 1A), that are packaged with the candicidin complex (Fig 1B,C, Fig S1A)^30^. After further concentration and extension of our bioactivity guided fractionation, we identified a second zone of weak antifungal activity corresponding to a cluster of peaks with spectra matching the antimycin A complex (Fig 1B,D, Fig S1A). Both metabolite complexes copurified in the same EV fraction after size exclusion chromatography and subsequent density gradient centrifugation (Fig S1B).

**Fig 1.**
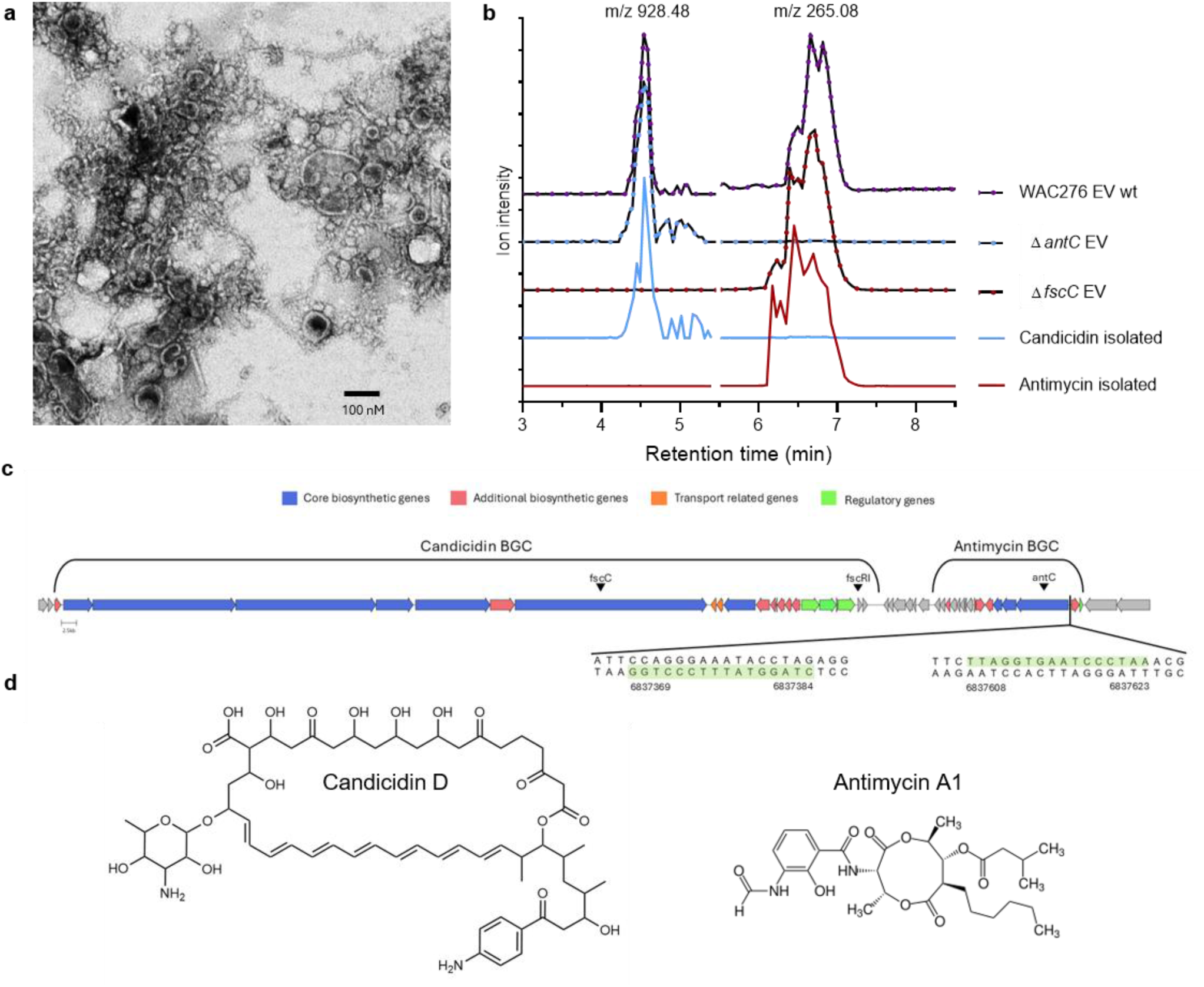
Extracellular vesicles from Streptomyces WAC276 carry candicidins and antimycins. **a**) Extracellular vesicles from WAC276 culture supernatant were concentrated and washed using 100 kDa ultrafiltration membranes and imaged by negative stain TEM (black bar 100 nM). **b)** Mass chromatogram traces for candicidins (extracted ion chromatogram of common fragment ion m/z 928.483 ±0.005, retention time 3-5.5 min), and antimycin A (extracted ion chromatogram of common fragment ion m/z 265.083 ±0.005, retention time 5.5-8.5 min). Samples are extracted EVs (black lines) from wild-type WAC276 (purified by size exclusion chromatography and subsequent density gradient centrifugation, purple dots), EVs from a strain with disrupted antimycin production (Δ*antC* EV, blue dots) or candicidin production (Δ*fscC* EV, red dots), and purified candicidins (blue line) and antimycin A (red line) from WAC276 culture supernatant. Traces are offset and have highest peak set to 100% intensity after normalization of each compound to wild-type EV m/z peak height. **c)** Biosynthetic region in WAC276 genome with >95% similarity to the reference clusters for candicidin and antimycin A, with genes coloured by predicted function. Black arrows highlight *fscC* – gene targeted for candicidin disruption, *antC* – gene targeted for antimycin disruption, and *fscR1* – regulator for polyene clusters. Green highlighted sequences show *fscR1* binding sites in antimycin cluster. **d)** Chemical structures for candicidin D and antimycin A1, the major analogues of each complex.

To examine the gene architecture responsible for this coproduced cocktail, we sequenced the genome of WAC276 and confirmed the presence of BGCs with high similarity (95% and 100% identity respectively) to the MIBig clusters for candicidin and antimycin (antismash v8.0)^54,55^. In WAC276 these BGCs are co-localized as a supercluster. The cluster encodes the candicidin regulator *fscC*R1, and FscCR1binding sites are present in the antimycin BGC (Fig 1C). This is the same supercluster architecture previously reported for coordinate regulation in strain *S. albidoffavus* S4^46,56^. To confirm these clusters were responsible for the EV metabolites causing the observed zones of antifungal activity, we generated mutant WAC276 strains bearing disruptions in the *fscC* PKS gene in the candicidin cluster and the *antC* NRPS biosynthetic gene in the antimycin cluster^34,57^. LC-MS/MS confirmed that the EVs from the Δ*fscC* strain no longer contained candicidins, but still contained antimycin A (Fig 1B, Table S1). EVs from the Δ*antC* strain no longer contained antimycins but still contained candicidins (Fig 1B, Table S1). In parallel, we purified the candicidin complex and the antimycin complex from EV-enriched WAC276 culture supernatants (Fig 1B, FigS1C-E). Using quantitative UV-Vis and LC-MS/MS the concentrations of candicidins and antimycins were determined in each of the wild type and mutant EV preps (Table S1). These data rigorously confirms WAC276 releases a population of EVs that contain candicidin and antimycin, a supercluster cocktail.

### The candicidin and antimycin supercluster occurs in two distantly-related clades of *Streptomyces*

In parallel with this biochemical investigation, we conducted a bioinformatic analysis to examine the phylogenetic and biosynthetic context of the candicidin and antimycin combination. We used the antiSMASH database^58^ of BGC predictions to identify all genomes that contained the candicidin and/or antimycin BGC and then filtered by species identification which yielded 28 distinct isolates with high-quality genomes for comparison (Table S2). Phylogenetic comparison of these strains along with our isolate WAC276 revealed that candicidin and antimycin BGCs are spread across a wide breadth of evolutionary distance (1.62 substitutions per site) in agreement with previous reports on the individual clusters^59–61^(Fig 2A). The antimycin BGC was the more widespread cluster, in 27 of the 29 strains. The candicidin BGC was found in 11 strains and co-occurs with the antimycin BGC in 9 of 11 strains (Fig 2A).

**Fig 2.**
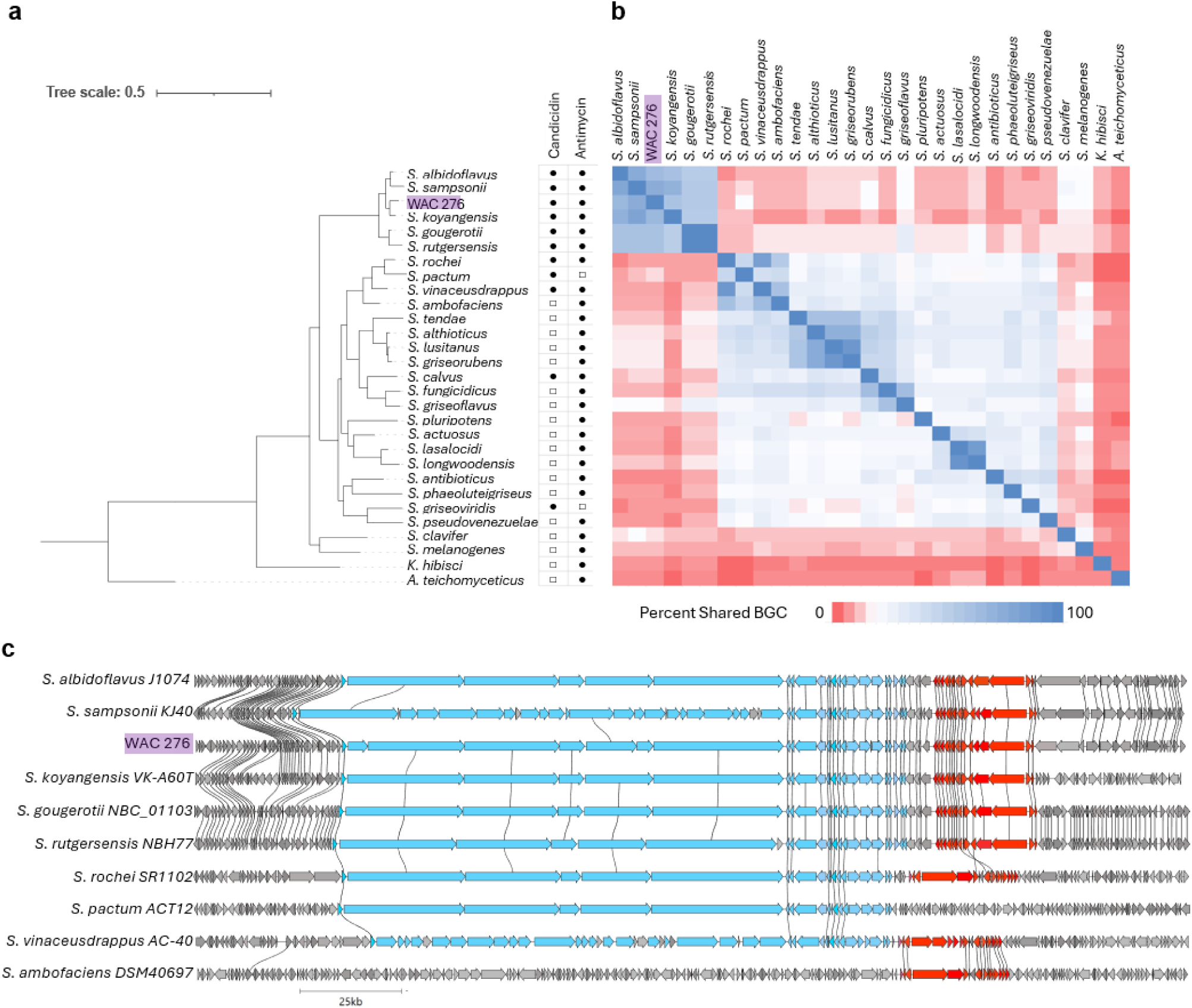
Occurrence and co-occurrence of candicidin and antimycin biosynthetic gene clusters. **a)** Phylogenetic tree of WAC276 and 28 strains found to contain either a candicidin or an antimycin BGC from the antiSMASH database, constructed using comparison of conserved markers through PhyloPhlAn 3.0^62^, branch length indicates substitutions per site. Presence of BGC indicated by black circle, absence by white square. WAC276 is highlighted in purple. **b)** Pairwise biosynthetic similarity of all strains, calculated as (number of shared BGCs in strain X and Y) / (total BGCs in strain X), X=row, Y=column position. Shared BGCs determined by BiG-SCAPE (cutoff 0.5). **c)** Clinker alignment of the candicidin and/or antimycin BGCs in indicated species. Blue is the candicidin BGC, red is the antimycin BGC, connecting line indicates ≥80% sequence identity.

To compare the biosynthetic relatedness of these 29 strains, we used a networking approach to cluster biosynthetic regions by gene similarity (BiG-SCAPE, cutoff 0.5, Fig S2, Table S2) and created a pairwise heatmap depicting the percent of shared BGCs between every strain (Fig 2B). This revealed that similar to phylogeny, the antimycin and candicidin BGC are found both alone and together in very disparate biosynthetic contexts. For example, WAC276 and *Streptomyces rochei* SR1102 both have candicidin and antimycin, but share <10% of their BGC repertoire. Six of the strains that contained both BGCs, including *Streptomyces albidoflavus* J1074/R2 and WAC276, were in a very closely related phylogenetic clade (maximum pairwise patristic distance of 0.2); however, they varied in as much as 50% of their biosynthetic repertoire. WAC276 had the highest biosynthetic overlap with *Streptomyces sampsonii* KJ40, sharing 75% of BGCs (Fig 2B).

We then examined the location of the candicidin and antimycin BGCs and found in 8 of 9 strains carrying both candicidin and antimycin that the BGCs are co-localized as a supercluster (Fig 2C). *Streptomyces calvus* did not have co-localized BGCs; however, it was unclear if the candicidin BGC in *S. calvus* DSM 41452 is functional as the candicidin BGC was interrupted and spread across two locations in the genome (Fig S2). In WAC276 and the 5 closely-related strains in the *S. albidoflavus* clade the arrangement of the candicidin and antimycin BGC subcomponents was virtually identical (Fig 2C). The other two strains carrying candicidin and antimycin as a supercluster were *S. rochei* SR1102 and *Streptomyces vinaceusdrappus* AC-40, with 86% biosynthetic similarity to each other, but less than 15% similarity to the *S. albidoflavus* clade. Interestingly they formed a clade with *Streptomyces pactum* ACT12, which contained a candicidin but no antimycin BGC, and *Streptomyces ambofaciens* DSM 40697, with an antimycin but no candicidin BGC (Fig 2A,C). In *S. rochei* and *S. vinaceusdrappus* the antimycin BGC is oriented in the opposite direction to the *S. albidoflavus* supercluster (Fig 2C). Moreover the antimycin BGC is in the intermediate form and lacks the *FscCR1* binding site^46,59^. This finding of co-localized candicidin and antimycin BGCs in two very diverse clades of *Streptomyces* and in varying strain-level biosynthetic repertoires is consistent with the hypothesis of selective pressures for supercluster architecture^18,20^. We next investigated the combination activity of the metabolites along with the impact of vesicle packaging.

### WAC276 EVs have strong antibiofilm activity requiring both candicidins and antimycins

We examined the contribution of candicidin and antimycin to EV bioactivity by comparing activity of wild-type EVs, EVs lacking metabolites (*ΔfscC* for loss of candicidin, *ΔantC* for loss of antimycin), and the purified metabolites (Fig 3, Fig S3)^34^. We found that candicidin is the dominant antifungal against planktonic growth of yeast (*Candida albicans, Cryptococcus neoformans* Fig 3, FigS3, *Saccharomyces cerevisiae* and other *Candida* sp. Fig S3), with antimycin contributing small gains in activity in non-fermentable carbon sources. Candicidins and other polyene antifungals disrupt cell membranes through their interaction with ergosterol^63^, while antimycin As inhibit cytochrome bc1, complex III of the mitochondrial electron transport chain, blocking aerobic respiration^49^. As expected, EVs containing candicidins (wt and *ΔantC*) as well as purified candicidins alone had potent activity on planktonic growth of yeast (Fig 3A-C, Fig S3A-D). In non-fermentable glycerol as a carbon source, EVs containing both antimycin and candicidin were 2-fold more potent than EVs lacking antimycin (Fig 3A wt vs *ΔantC*). However, in fermentable glucose loss of antimycin in EVs or addition of purified antimycin to candicidin in combination did not change potency (Fig 3A-C). Similarly purified antimycin, or EVs containing antimycin but lacking candicidin (*ΔfscC*), had detectable activity on glycerol, but weak and/or incomplete inhibition in glucose (Fig 3A-C, Fig S3A-C). This likely reflects the fact that during growth on glucose, yeast can bypass the mitochondrial electron transport chain and rely on fermentation rendering antimycin ineffective. To assess the impact of vesicle packaging we compared wild-type WAC276 EVs to a matched ratio of purified candicidin and antimycin, and found the EVs were consistently 2-4x more potent (Fig 3B,C, Fig S3A-C).

**Figure 3.**
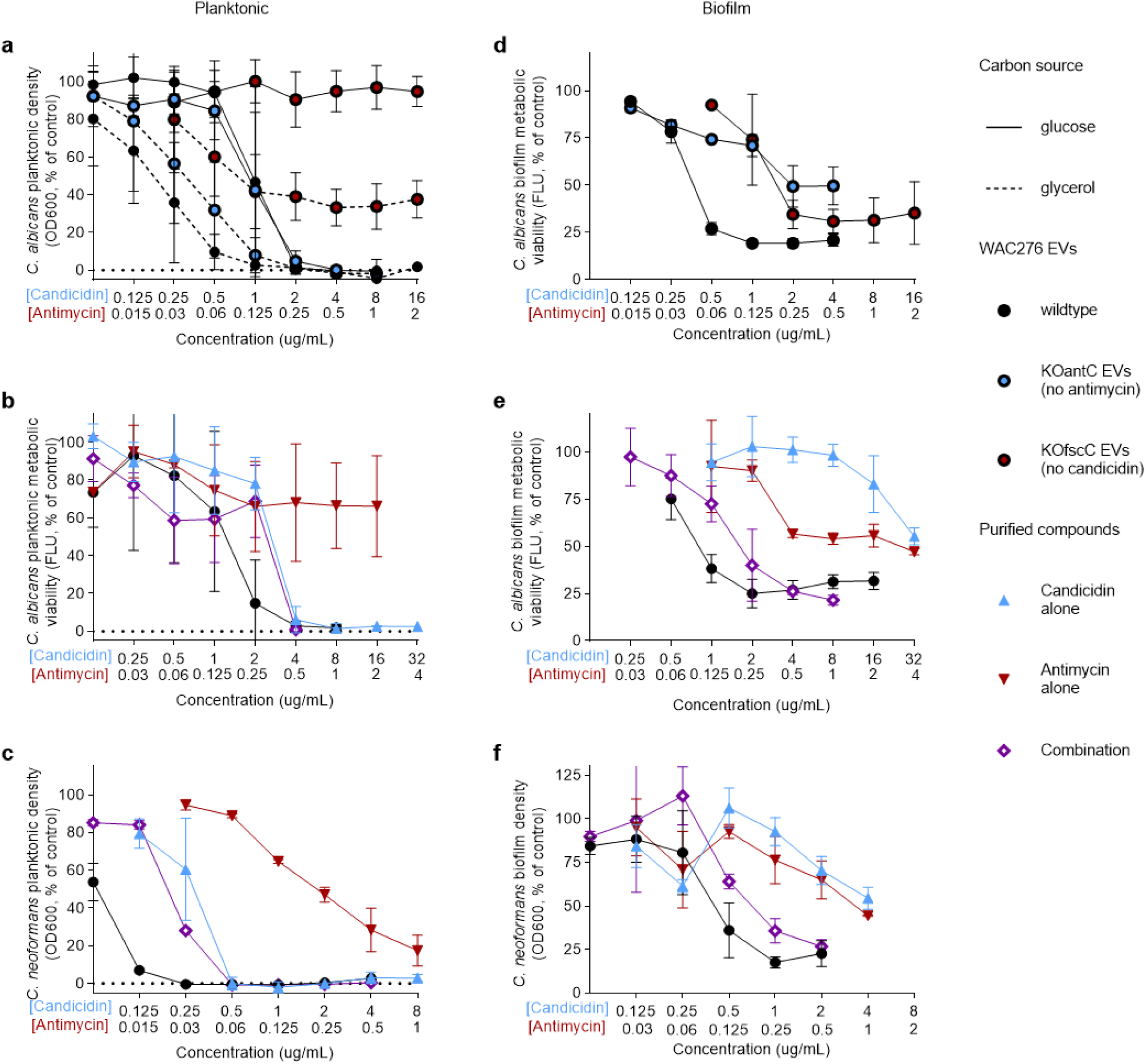
Contribution of candicidin and antimycin to WAC276 EV antifungal activity against planktonic cells and biofilms. **a, b, c,** Planktonic inhibition, and **d, e, f,** biofilm inhibition of *C. albicans* (a, b, d, e) or *C. neoformans* (c, f). WAC276 wildtype EVs (black circles) containing both candicidin and antimycin, were compared to EVs with candicidin but no antimycin (blue circles) or EVs with antimycin but not candicidin (red circles) (a, d), or to purified candicidin (blue triangles), antimycin (red triangles), or a matched ratio of candicidin and antimycin (purple diamonds) (b, c, e, f). Inhibition was tested after overnight incubation at 37°C in medium with either glucose (solid lines) or glycerol (dashed lines) as a carbon source. The graphs show the metabolic viability or cell density of treated cells compared to untreated controls, and the relevant candicidin or antimycin concentration can be read on the X axis, candicidin concentrations top row and antimycin concentrations second row. Symbols are means, error bars are SD, for planktonic assays of biological replicates, *C. albicans* (n≥3), and *C. neoformans* (n≥2), and for biofilms of technical replicates (n≥3) of a representative assay from at least biological duplicate.

We then examined activity against mature biofilms, a critical ecological and clinical mode of growth for many microbes including yeast. We found the activity of candicidin in combination with antimycin was strikingly different to that of planktonic growth. Wildtype WAC276 EVs and the purified combination of candicidin and antimycin had excellent efficacy against mature fungal biofilms in glucose, maintaining similar potencies to planktonic inhibition (Fig 3D-F, Fig S3E,F). In contrast, loss of either antifungal in the EV substantially reduced antibiofilm potency and efficacy (Fig 3D), and EVs lacking both metabolites (from strains with double disruptions, *ΔfscCΔantC*) were 100x less active (Fig S3G); providing strong evidence that the combination of these supercluster derived metabolites is a vital component of the antibiofilm efficacy of the EVs. Similarly, mature *C. albicans* or *C. neoformans* biofilms tolerated high concentrations of purified candicidins, where concentrations 8x above planktonic MIC achieved just 50% inhibition (Fig 3D,E). Interestingly antimycins had improved activity against *C. albicans* biofilm viability compared to planktonic inhibition in glucose; however, as a single compound antimycins also had a limited maximum efficacy of only 50% (Fig3 B,D). Notably the combination of candicidin and antimycin had excellent potency against the biofilms, and a greatly improved maximum efficacy (Emax) of 80% compared to the individual compounds (Fig 3D,E). Again, WAC276 EVs were consistently 2-fold more potent than the matched purified combination (Fig 3D,E, Fig S3E). Together this data shows that candicidin and antimycin display a contingent coordinated activity in combination against biofilm growth of fungi, and packaging in EVs enhances antifungal activity.

### Simultaneous disruption of ergosterol and the electron transport confers synergy against *C. albicans* biofilms

To assess the extent of synergy of candicidins and antimycins against mature *C. albicans* biofilms, we tested candicidins and antimycins at various ratios, ranging from their individual effective concentrations (64:1) to ratios we observed in our WAC276 EVs (8:1, 4:1) (Fig 4 A,B). All ratios were highly synergistic as demonstrated by a concave isobologram of the combined dose required to inhibit 50% biofilm viability (Fig 3C). Although the candicidin and antimycin combination was not cidal against mature biofilms (Fig S4A), we also evaluated biofilm prevention conditions and observed synergistic biofilm prevention, and once again an even greater potency of the WAC276 EVs (Fig S4B). Hyphae formation is an essential component of fungal biofilm formation for yeast such as *C. albicans*. We examined the impact of individual metabolites, the combination, and WAC276 EVs on *C. albicans* hyphenation in high glucose at 37°C, our biofilm condition (Fig 4D, FigS4C). At sub-MIC concentrations, antimycin A did not appear to reduce filament length relative to untreated controls, whereas candicidin reduced hyphal elongation. The combination of candicidin and antimycin and the WAC276 EVs displayed both a reduction in hyphal elongation and replication inhibition (Fig 4D), similar to the enhanced combination effect against biofilm growth.

**Figure 4.**
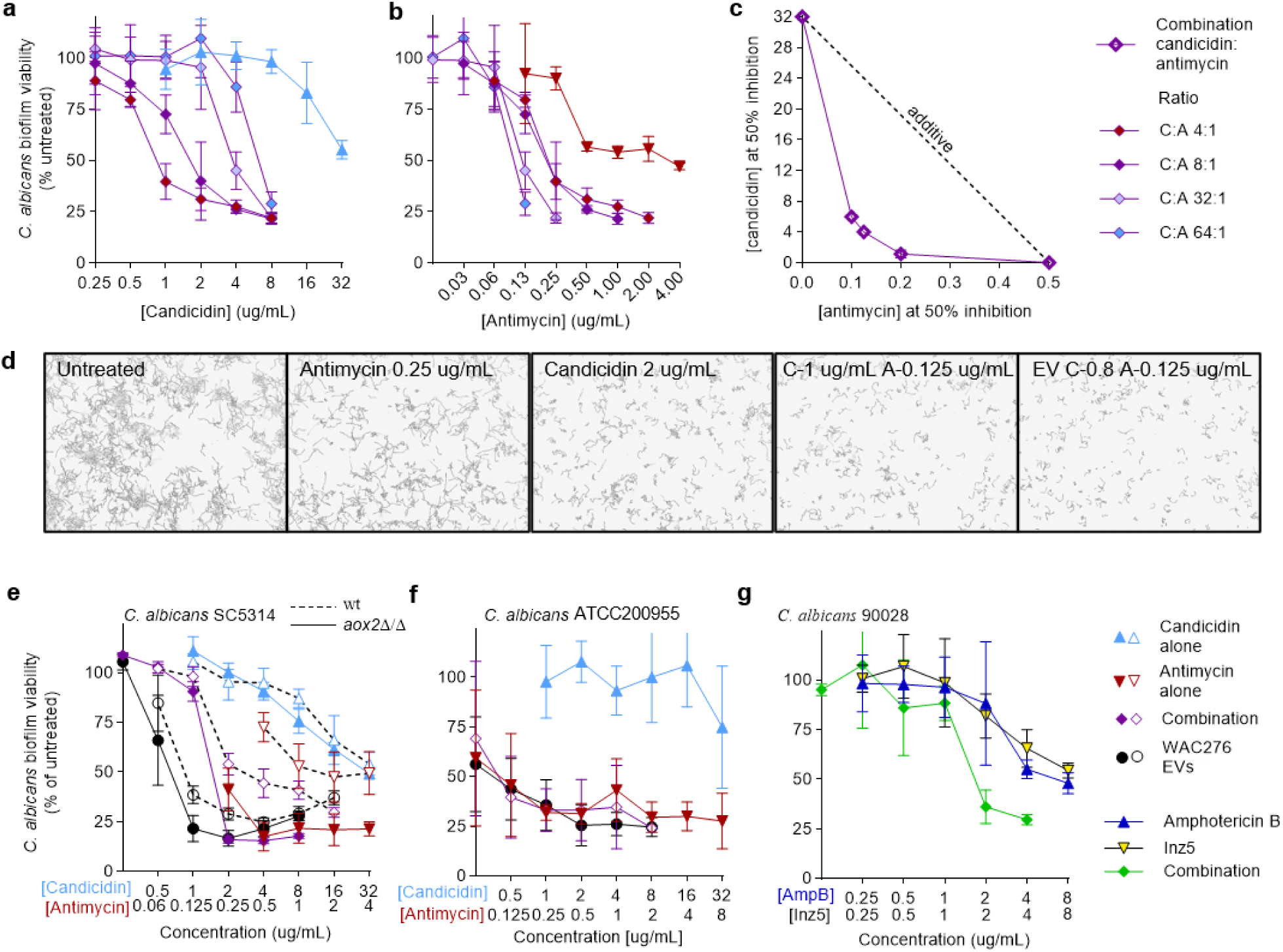
Combined sensitivity to ergosterol and electron transport chain disruption in *C. albicans* biofilms. **a-c)** Mature *C. albicans* biofilm sensitivity to candicidin (alone, blue triangles) and antimycin (alone, red triangles), and combined in various combination ratios (purple diamonds, blue fill 64:1, light purple fill 32:1, solid purple 8:1, red fill 4:1). Plotted for candicidin concentration **(a)**, or antimycin concentration **(b)**, and the doses in combination required to reduce viability to 50% of untreated control are plotted on an isobologram in **c)**. **d)** Biofilm formation conditions of high glucose at 37°C shown after 6 h. Untreated *C. albicans* form lots of hyphae in dense clumps, whereas treatment with antimycin or candicidin (at 0.5xMIC) alone slows extent of hyphenation and clumps, and with even lower concentrations the combination or treatment with WAC276 EVs is more effective still. **e)** Biofilms of *C. albicans* with a mutation in alternative oxidase (*aox2*Δ/Δ, black lines) are more sensitive to antimycin (red triangles), combination (purple diamonds) and WAC276 EVs (black circles) treatment than wild-type biofilms (dashed lines) treated in parallel. **f)** *C. albicans* biofilms with a clinical isolate (ATCC90028) harbouring a mutation in the ergosterol biosynthesis gene (*ERG2*) are tolerant to candicidin alone (blue triangles), but hypersensitive to antimycin alone (red triangles), with the combination of candicidins and antimycins (purple diamonds) and WAC276 EVs (black circles) displaying no better inhibition than the antimycins alone. **g)** The clinically used polyene amphotericin B (blue triangles) and the indazole fungal specific cytochrome bc1 inhibitor Inz-5 (yellow triangles) have limited biofilm efficacy alone, with enhanced activity in combination (green diamonds) against wild-type *C. albicans*. Symbols are means, error bars SD, of technical replicates (n≥3) of a representative biofilm assay from at least biological duplicate.

To assess the mechanistic role of candicidin and/or antimycin in the synergistic antibiofilm activity, we grew biofilms of a *C. albicans* strain with a mutation in alternative oxidase (*aox2*Δ/Δ) forcing reliance on the ETC (Fig 4E), and biofilms of a *C. albicans* clinical isolate with a mutation in the ergosterol biosynthesis gene *ERG2* that displays resistance to amphotericin (ATCC200955)^64,65^. The *aox2*Δ/Δ biofilms had increased sensitivity to antimycins, the combination, and WAC276 EVs compared to the parent *C. albicans* strain (SC5314) (Fig 4E), consistent with ETC inhibition contributing to coordinated activity. Antimycin had improved maximum efficacy against the *aox2*Δ/Δ biofilms, which suggests that the tolerance to antimycins of wild-type biofilms is, at least in part, due to substantial anaerobic respiration. The ATCC200955 biofilms were highly tolerant to candicidin alone (Fig 4F). Interestingly, there was no longer combination synergy or enhanced potency of EVs compared to purified compounds against the clinical isolate mutant (Fig 4F).

Notably however, biofilms of the polyene-resistant clinical isolate were extremely sensitive to antimycin A (Fig 4F). If this is a result of ergosterol pathway disruption, this would be an interesting collateral sensitivity effect. This hypersensitivity to antimycin A resulted in the combination and the EVs maintaining strong potency against the candicidin-resistant clinical isolate compared to the wild-type strain (Fig 3E, 4E-F).

To determine whether enhanced antibiofilm activity could occur with other ETC inhibitors and ergosterol inhibitors, we tested the indazole fungal-specific cytochrome bc1 inhibitor Inz-5^66^ alone or in combination with the clinical fungicidal polyene amphotericin B. While modest, we found that the combination of Inz5 with amphotericin B did indeed result in increased potency and greater combined maximum efficacy, similar to combinations of candicidin and antimycin (Fig 4G). Together, this data supports a mechanistic interplay between ergosterol disruption and electron transport chain inhibition devastating to *C. albicans* biofilms.

### EVs maintain concentrations and ratios of candicidins and antimycins for delivery

One critical question is the impact of EV packaging of two distinct compounds on their diffusion and delivery^67^. This is important to combination activity because compounds having differing diffusion properties might not arrive at target cells with sufficient concentration to realize their synergistic effect. It seems plausible therefore, that vesicle-packaging could enhance the benefits of co-production from a BGC supercluster.

To explore relative diffusion, we compared the zones of inhibition created by the purified candicidins and antimycins with those of WAC276 EVs on agar plates. The zone of inhibition created by an antimicrobial on an agar lawn of a target microbe relies on diffusion of the antimicrobial outwards from the spot of application^68^. Slow diffusion thus limits the increase in zone achieved with increased application of antimicrobial. Conversely, fast diffusion enables a large zone with high antimicrobial application, however, with limited amounts of antimicrobial fast diffusion rapidly dilutes the antimicrobial below inhibitory concentrations and thus the zone of inhibition is very responsive to amount applied^68^. Serial dilutions of purified antimycins and candicidins were spotted on agar plates seeded with *C. albicans* and *C. neoformans* and compared to intact vesicle preparations. High concentrations of purified antimycins created large zones of incomplete inhibition, and the zones shrank dramatically as the concentration decreased, indicating rapid diffusion of these compounds in the agar (Fig 5A,B). Purified candicidins created smaller zones of complete inhibition with only small changes in diameter as concentrations decreased, indicating limited diffusion in the agar. Vesicles, containing both candicidins and antimycins, generated zones of inhibition that shrank with a slope in between that of the purified compounds, consistent with the possibility that vesicles keep the candicidins and antimycins dually localized and change diffusion properties (Fig 5A,B). This dual localization would contribute to the greater inhibition achieved with lower minimum amounts of compounds observed for vesicles compared to purified compound combinations (Fig 5A,B).

**Figure 5.**
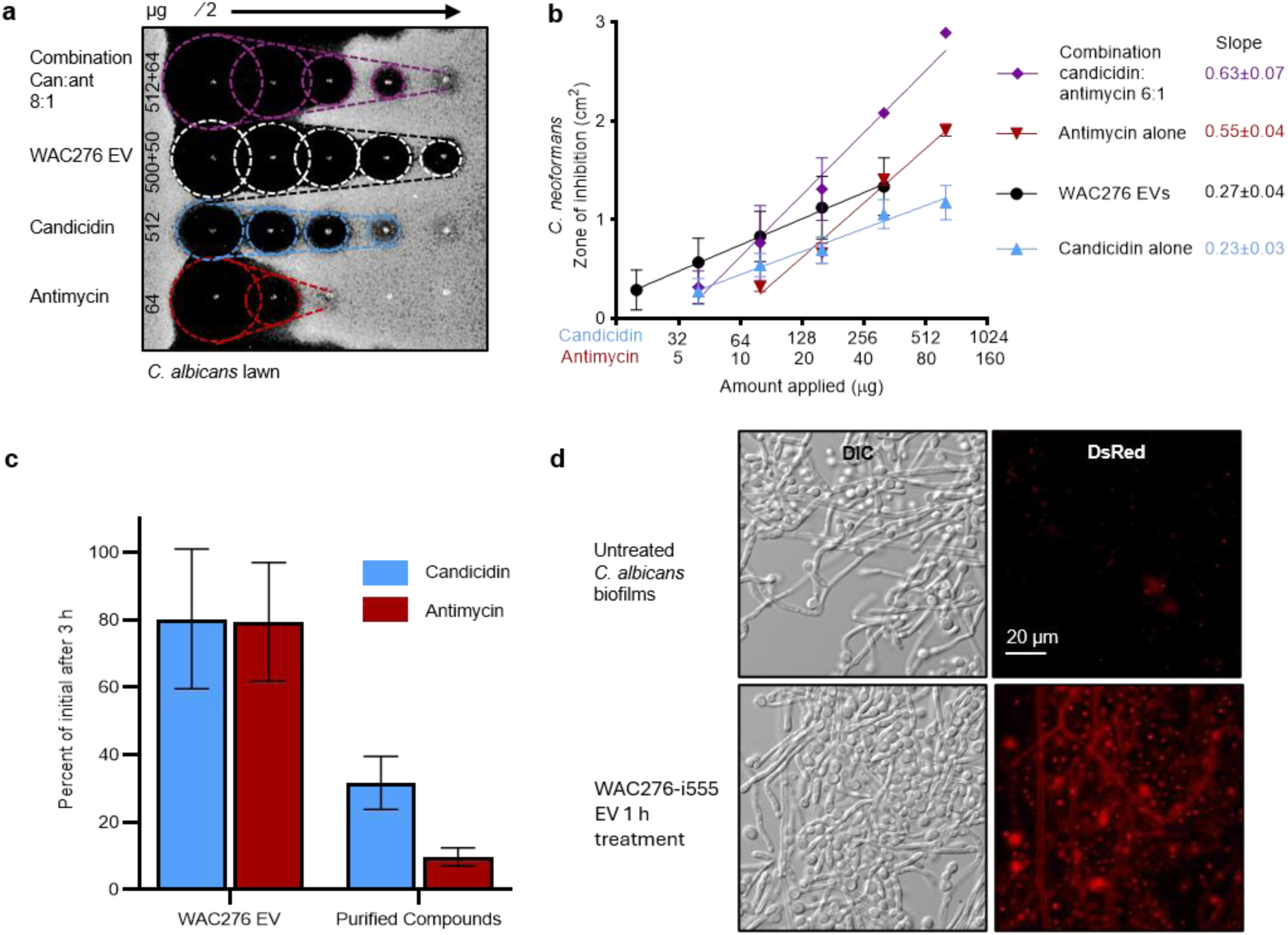
EV packaging preserves candicidin and antimycin ratios and enables delivery through biofilm. **a)** Change in zones of inhibition on serial dilution of purified compounds compared to WAC276 EVs against lawns of *C. albicans* on YPglycerol agar reflecting potency and diffusion from the centre spot of application. Zones and slope highlighted: combination of purified candicidin and antimycin, purple; wild-type WAC276 EV, white and black; candicidin, blue; antimycin, red. **b**) Quantification of areas of inhibition zones on *C. neoformans* compared to amount applied for purified individual compounds alone (candicidin – blue, antimycin -red), and in combination (purple), and WAC276 EVs (black), mean and SD, n=3 biological replicate. Slope±SD is linear regression of area/log_2_(candicidin), GraphPad Prism v11. **c**) Percent of candicidin (blue) and antimycin extractable from supernatant of sonicated mature *C. albicans* biofilms after 3 h incubation compared to initial treatment, mean and SD of n≥3 biological replicate. **d**) DIC and DsRed fluorescence images of mature *C. albicans* biofilms untreated or treated for 1 h with WAC276 EVs labelled with i555-SE, then washed, and scraped onto a microscopy slide.

Diffusion and delivery are a particularly important challenge in the context of biofilms. One mechanism of biofilm tolerance is the adsorption of antimicrobial compounds to the biofilm matrix and/or restricted diffusion. In both cases, the result is a reduced load of compound reaching the cell surface. To examine loss of candicidin and antimycin in the presence of biofilm matrix, we used LC-MS/MS to quantify the amounts of each metabolite complex remaining in the supernatant after 3 h incubation with our mature *C. albicans* biofilms. The purified compounds experienced dramatic loss to the biofilm with 70% of candicidin and 90% of antimycin A loss in 3 h, but we found that WAC276 EVs protected against that loss with 80% of each complex remaining (Fig 5C). Importantly, this also means EV packaging maintains the relative ratio of candicidins and antimycins through the matrix (Fig 5C, Fig S5). To investigate the potential of the WAC276 EVs to achieve delivery of cargo through the fungal biofilms, EVs were covalently labeled on external free amines with a small fluorescent tag (Alexa fluor 555). These EVs were added to mature *C. albicans* biofilms in rich medium (YPD), then biofilms were washed thoroughly and scraped onto slides for imaging. There was rapid acquisition of fluorescence into both hyphae and into cell bodies (Fig 5D), visible as early as 15 min in 40% of the fields of view across the slide, and visible in 100% of the fields of view by 3 h. The fluorescence in cell bodies appeared to be in endocytic compartments, consistent with the rapid endocytic turnover of yeast cell membranes^69^. The clear membrane labeling of the hyphae suggests that the WAC276 EVs achieve membrane fusion with the fungal cell membrane, and most importantly this data suggests EVs are able to diffuse effectively through the biofilm.

This ability of the EVs to ensure a matched co-delivery of the candicidin and antimycin cargo into cells across space and time is likely to be a critical feature of their enhanced potency compared to purified compounds (Fig 3, 4). It is possible that unknown components of the EVs further enhance efficiency, however, it is clear that co-delivery is advantageous for antifungal efficacy (Fig 5A-C). We suggest that this is a significant evolutionary advantage to the packaging of this combination into extracellular vesicles.

## DISCUSSION

The coordinated biology of microbial specialized metabolites is a frontier that provides new insights into the complexity of microbial interactions and the selective pressures acting on metabolite genetics and regulation. It also reveals novel bioactivities not possible with single molecules. We find an EV-packaged supercluster-encoded combination of metabolites that exhibit coordinated synergistic activity against fungal biofilms (Fig 1-4). The occurrence of this combination in diverse phylogenetic and biosynthetic backgrounds, and in 9 out of 11 *Streptomyces* species encoding candicidin, is consistent with maintenance of the combination through positive selection (Fig 2). The combination activity of the ergosterol targeting polyene candicidin and the ETC inhibitor antimycin A is sensitive to target genetic modifications, implicating a mechanistic vulnerability of fungal biofilms exploited by this coordinated combination (Fig 4). Notably packaging in EVs provides a protective and a co-delivery benefit and enhances antifungal activity (Fig 3,5). Fungal biofilm growth is problematic in many clinical and industrial settings. Inspired by this natural antibiofilm EV cocktail, further exploration of co-encapsulated polyenes and ETC inhibitors may be beneficial given the widespread use of ergosterol-targeting polyenes derived from *Streptomyces* in agriculture, food preservation, and medicine.

We find supercluster architecture of the candicidin and antimycin BGCs in both the *Streptomyces albidoffavus* and the *Streptomyces rochei* clade (Fig 1,2). In the six *S. albidoflavus* clade species, including our isolate WAC276, the supercluster arrangements are virtually identical (Fig 2C), and based on previous studies are likely to be co-regulated by the polyene regulator FscR1^46^. In *S. rochei* and *S. vinaceusdrappus,* the antimycin A BGC is in the opposite orientation to the *S. albidoflavus* supercluster, and lacks evident FscR1 binding sites, leading to an open question for future study of whether co-ordinated production of candicidin and antimycin occurs in these strains. The candicidin BGC occurs rarely without antimycin – in our analysis we found this only in *S. pactum* and *Streptomyces griseoflavus*. Co-localization in a supercluster across distantly related species has been observed for other known examples of synergistic combinations^18,19^, and our data with candicidins and antimycins adds to the evidence that synergy is a driving force for the evolution of supercluster architecture^18^. Candicidin and antimycin potentiation is not evident under the commonly tested laboratory conditions of planktonic growth and abundant glucose. Instead, we observe synergy against fungal biofilms, and additivity against yeast grown in non-fermentable carbon sources – both common conditions for microbial growth in the environment. This reconciles previous disparate observations on the combination activity of candicidins and antimycins^46,70,71^. A conditional benefit of producing a second molecule emphasizes the contingency aspect frequently observed with coordinate bacterial specialized metabolite production^18,23^.

We identify WAC276 as an isolate of *Streptomyces albidoflavus* where candicidins and antimycins are both packaged in EVs (Fig 1). Candicidins have been associated with secreted lipid particulate/EVs in several strains^30,34,72^, suggesting this may be a ubiquitous secretion mechanism. To our knowledge antimycins have not previously been identified as vesicle cargo^34^, and given the wide breadth of antimycin producers, it would be an interesting future question to examine the frequency of secretion in EVs. The WAC276 EVs have enhanced activity compared to purified candicidins and antimycins, consistently demonstrating 2-4x greater potency than a matched combination in planktonic and biofilm assays (Fig 3, 4). Microbial growth in biofilms is ubiquitous in the environment and infections^73^. Biofilm tolerance to antimicrobials is multifactorial, with slowed metabolism, growth, altered transcriptional programs, as well as delivery through the extracellular matrix representing factors that can reduce antimicrobial efficacy^52,74–77^. Notably, we find an example of a combination with mechanistic synergy against fungal biofilms involving ergosterol disruption and ETC inhibition, as well as packaging in EVs, which improves delivery properties. Although antimycin A does not have translational potential for humans due to affinity for human cytochrome bc1, for proof-of-concept we tested the combination of amphotericin B and a fungal selective cytochrome bc1 inhibitor, Inz5^66^, and also observed potentiation against biofilms (Fig 4G). Our results add to the accumulation of evidence of a vulnerability of fungi to both membrane disruption and ETC inhibition^78^, for example in reports of synergy between Inz5 and azoles, and lansoprazole and amphotericin B^64,79^. Our data also supports the vulnerability of fungal biofilms to respiration inhibition^78^; however, our results of 50% maximum inhibition of antimycin or Inz5 alone (Fig 3,4) would caution against the sole use of respiration inhibitors. Interestingly, in an amphotericin B-resistant clinical isolate of *C. albicans* with mutations in the ergosterol biosynthesis gene *ERG2*^64^, we observed a hypersensitivity to antimycin under biofilm conditions (Fig 4F), allowing maintenance of maximal inhibition by the combination of candicidin and antimycin at potencies similar to wild-type *C. albicans*. Biofilms are of particular concern as incubators and reservoirs for acquired antimicrobial resistance. Candicidin and antimycin are a naturally selected for specialized metabolite combination able to counter both intrinsic and acquired antimicrobial resistance in biofilms.

One challenge for coordinated activity of combinations is achieving favourable exposure kinetics to the substituent molecules. EVs are able to provide an intercellular transport mechanism for cocktails of very disparate molecules^80–82^. In our data, EV packaging had an altered diffusion pattern of antifungal inhibition on agar plates compared to isolated compounds^68^. EVs had reduced diffusion compared to isolated antimycins and increased diffusion compared to isolated candicidins (Fig 5A-C), which therefore would maintain more constant ratios than the purified compounds in combination, and increase the area of combined effect when amounts of the metabolites are limited^68^. We also found EV packaging protected against the loss of candicidin and antimycin in biofilm treatment conditions, importantly maintaining the initial ratio of compounds (Fig 5D). Furthermore EVs were able to fuse into biofilm cell membranes (Fig 5E). Previous studies of the polyene amphotericin B have shown that amphotericin B binds readily to the β-glucans of the *C. albicans* biofilm matrix^83^, whereas liposomal formulations of amphotericin B reduce this loss and show enhanced biofilm efficacy^84^. We note with interest this convergence of the benefits of the empirically developed clinical liposomal formulation of the polyene amphotericin B (a specialized metabolite of *Streptomyces nodusus*) with our findings on the naturally released bacterial EV packaging of the polyene candicidin. The ability of the EVs to keep candicidins and antimycins together and to deliver cargo into cells may explain the increased potency and efficacy of the EVs compared to purified compounds, although our results do not rule out the contribution of other components of the EVs which we plan to investigate in future studies. We note that the modified diffusion and delivery properties of specialized metabolites in EVs may also provide a contingency benefit to producing cells, enhancing bioactivity under certain environmental conditions where diffusion and delivery are limiting^85–88^, including in biofilm environments^89–93^.

In conclusion, our study provides further evidence that the combination specialized metabolite products of biosynthetic superclusters can exhibit coordinate activity, particularly in the context of ecologically relevant conditions such as carbon source and biofilm growth. We find candicidins and antimycins in combination have contingent synergy against biofilms, countering the intrinsic resistance of biofilms of *C. neoformans* and *C. albicans* to the individual antifungals. We find an isolate of *Streptomyces albidoffavus* (WAC276) that packages this supercluster combination into EVs, to our knowledge the first observation of an EV packaging of supercluster combination. The EVs protect and maintain the concentrations of the two antifungals, and achieve delivery of cargo into cells under biofilm conditions. Our study provides evidence that bacterial EVs can act as delivery vehicles for cocktails of specialized metabolites, and that EVs can have useful delivery properties in microbial biofilms.

## METHODS

### Strains, Culture Conditions, And Compounds

*Streptomyces* isolate WAC276 and derivatives were maintained as aliquoted glycerol spore stocks, and cultures were directly inoculated from spore stocks at final 5x10^6^CFU/mL into Maltose Yeast Extract Malt (MYM) medium (components from Becton Dickinson) with added trace elements^94^. Yeast were maintained as glycerol stocks, then streaked onto Yeast Peptone Dextrose (YPD) agar plates and cultured overnight at 30°C for single colony isolation. Overnight cultures were grown from single colonies in YPD with shaking at 30°C. Cultures were then diluted for biofilm assays or planktonic assays at 37°C. Yeast strains: *C. neoformans* H99, *C. albicans* ATCC90028 (ATCC), *C. albicans* SC5315, *C. albicans* aox2Δ/Δ^95^, *C. albicans* ATCC200955 (*erg2*, ATCC). Amphotericin B (Sigma-Aldrich, A4888), Inz-5 (MedChem Express, HY-121721), and purified compounds were stored as DMSO stocks at -80°C in multiple aliquots. Alexa Fluor 555 NHS ester (Invitrogen) was stored in DMSO at -20°C.

### Transmission Electron Microscopy

Culture supernatant from WAC276 was concentrated 10x and washed 100x with PBS on a 100kDa Amicon 500 uL filter (spun at 8000xg), and then incubated on negatively charged carbon grids (CF400CU50, EMS) for 60 s. Grids were washed three times with water, and stained with 2% uranyl acetate for 30 s. Grids were imaged using a Talos L120C transmission electron microscope (software TEM Imaging & Analysis v5.0 SP4).

### Construction Of *Streptomyces* WAC276 Mutant Strains

To disrupt *fscC* and *antC*, a gene disruption technique integrating an antibiotic selection marker into the gene by homologous recombination was used. A 1 kb region near the 5’ end of each gene was amplified from WAC276 genomic DNA (isolated by Dneasy Blood & Tissue kit, Qiagen) (Primers, Table S3) and cloned into the HindIII site of pOJ260^96^ to generate pOJ-fscC or pOJ-antC. Following sequence verification, pOJ-fscC/antC was introduced into *E. coli* ET12567/pUZ8002 and subsequently conjugated into WAC276.

Genomic DNA of apramycin-resistant exconjugants was isolated to confirm disruption of *fscC* or *antC* via PCR using primers targeting the integration site and verified by sequencing.

### Extracellular Vesicle Isolation

Seven-day old WAC276 liquid cultures were pelleted at 3000xg for 20 minutes. 40 ml of supernatant was collected from each pelleted culture and filtered using a 0.2 um PES membrane to remove larger aggregates and any remaining cells. The filtered supernatant was then concentrated (10kDa Amicon 15 ml) at 3500xg for 20 minutes repeatedly, until each initial 40 ml of supernatant was reduced to 1 ml. After pelleting any aggregates, the concentrated supernatant was applied to size exclusion chromatography (SEC) columns (qEV1-35nm, Izon) and eluted with PBS. After a void volume of 4.5 mL, three 1 mL EV-containing fractions (F1, F2, and F3) were collected and combined, and stored as 50µl aliquots at -80°C. Aliquots were single use, vesicles were thawed rapidly and spun at 5000xg for 5 minutes to pellet aggregates, then supernatant used immediately.

### Differential Gradient Centrifugation

To confirm candicidin and antimycin EVs co-purified, two batches of WAC276 wt EVs were further purified by differential gradient centrifugation following SEC. 1 mL of EVs was mixed with 2 mL of 60% iodixanol (Optiprep) and layered at the bottom of a step gradient of 2 mL each of 35% to 20% iodixanol in 5% steps (from bottom to top) in thin wall 13.2 mL tubes (Beckman Coulter 331372). The tubes were ultracentrifuged at 100,000x*g* for 16 h [4°C, SW41Ti rotor, Optima XPN-80 centrifuge, Beckman Coulter], and then 0.5 mL fractions were removed by pipette sequentially beginning from the top of the tubes. 25 uL of each fraction was then extracted with final 75% methanol and analyzed by LC-MS/MS.

### Metabolite Purification

WAC276 cultures were inoculated with 5x10^6^CFU/mL in MYM (8 flasks of 50 mL) and shaken at 220 rpm for 7 days at 30°C. Cells were removed by centrifugation (5000xg, 20 min), and supernatant concentrated on 10 kDa Amicon 15 mL ultrafiltration membranes from 350 mL to 80 mL. Concentrated supernatant was extracted overnight at 4°C with equal volume butanol. The butanol layer was dried completely (Genevac EZ-2 Elite), and resolubilized in 4 mL 75% methanol. After spinning down precipitate, sample was loaded onto two solidphase C18 columns (Waters Sep-Pak) and eluted with 20 mL of sequential water, 10% acetonitrile, 20% acetonitrile, 50% acetonitrile, 75% acetonitrile, 100% acetonitrile, and then 100% methanol. The 50% acetonitrile fractions contained the candicidins, and were dried and resolubilized in 500 uL of 50% acetonitrile, and then separated on an analytical PFP column (4.6x250mm, Phenomenex) by HPLC (Alliance) using a step gradient. Candicidin containing fractions were identified by the triple polyene absorbance peaks (Fig S1A) and collected, dried, weighed (yield ∼1mg from 400 mL), and dissolved in DMSO then aliquoted and stored at -80°C. The 75% and 100% acetonitrile fraction contained the antimycins. These fractions were dried, and resolubilized in 500 uL of 100% acetonitrile and separated on an analytical C18 column (4.6x250mm Luna C18 5 µm 100 Å Phenomenex). Fractions containing antimcyins were identified by absorbance at 320 nm (FigS1A), collected, dried, weighed (yield ∼1mg from 400 ml), dissolved in DMSO, aliquoted and stored at -80°C.

### LC/MS-MS Compound Quantification

EV aliquots stored at -80°C were thawed and spun at 5000xg for 5 min to remove aggregates. A series of EV extracts were prepared by spiking in either DMSO alone or an internal standard of the purified candicidin and antimycins at three concentrations, and then methanol added to final concentration 65%. Standard curves of purified candicidins and antimycins were prepared in the same solvent backgrounds of 25% PBS, 10% DMSO, 65% methanol, ranging from 1 ug/mL to 25 ug/mL. Samples were analyzed by LC-MS/MS - Acquity UPLC I-Class coupled to a Xevo G2-S QToF (Waters). 10 uL samples were separated on an Acquity BEH C18 column (1.7um, 2.1x50mm) at 40°C with 0.2 mL/min flow, beginning with 95% water + 0.1% formic acid (A), 5% acetonitrile + 0.1% formic acid (B), held for 1 min, then changing to 95% B over 5min, held at 95% B for 2 min, and returned to 95% water over 1 min and held for a final 1 min. PDA detector collected absorbance from 190 to 800 nm with a resolution of 1.2 nm. Ionization was ESI, positive mode, and MS spectra were collected over the whole run for 50-1200m/z with a scan time of 0.5s (corrected by lock mass). For antimycin quantitation, MSMS spectra of the common antimycin base peak 265.1 were collected from 5.7 to 7.5 min, with a collision energy of 14eV and scan time of 0.5s. To quantify candicidin, targetlynx software was used to determine peak area for 380+/-30 nm from the PDA function for retention time 4.2 to 5 min. EV samples were diluted so that peak area would be in the quantifiable range and candicidin concentration calculated from standard curves, with each sample quantified on at least two separate occasions (Table S1). To quantify antimycin As, targetlyx software was used to determine the peak area of m/z 237.08 +/- 0.01 in the MSMS m/z 265.1 function for each EV sample alone and with the three spiked internal standards of the purified antimycin, a linear regression fit to the curve, and the y-intercept and slope used to calculate the EV antimycin concentration, with each sample quantified on at least two separate occasions (Table S1).

### Bioinformatic Phylogeny And Biosynthetic Capacity

Strains used for bioinformatic analysis were found through the antiSMASH database v5.0^58^. The query function was used to find strains that had a KnownClusterBlast hit of the antimycin BGC only (BGC0000958), the candicidin BGC only (BGC0000034), or both antimycin and candicidin BGCs. BGCs with percent similarity <85% or found on a contig edge were excluded. Search results were further narrowed down to exclude unknown species and genomes containing more than 5 contigs, and then a representative strain was chosen for each species. To confirm the presence of either candicidin BGC or antimycin BGC, the BiG-SCAPE v2.0^97^ query was used including the MiBig clusters (BGC0000034 and BGC0000958, respectively) with a cutoff of 0.5. nBLAST against *fscC* (AAQ82564.1) or *antC* (WP_010639240.1) was used to further validate the presence/absence of the candicidin BGC or antimycin BGC, respectively. A cutoff of an E-value of 0, 80% identity, and 80% query coverage was used. This resulted in 28 strains remaining (Table S2).

Genomes were downloaded as RefSeq FASTA files from the National Center for Biotechnology Information and inputted into PhyloPhlAn v3.0^62^. The phylogenetic comparison aligned the phylophan database’s 400 universal marker genes found in the input genomes. Analyses were run with the diversity medium parameter and the default supermatrix_aa configuration file. The pipeline utilized DIAMOND (v0.9.19), MAFFT (v7.525), trimAl (v1.5), Fasttree (v2.2.0), and RAxML (v8.2.12). The resulting maximum-likelihood tree was visualized and rooted with respect to *Actinoplanes teichomyceticus* using iTOL v7^98^. Branch lengths indicate substitutions per site.

The BGCs of each strain were obtained from the antiSMASH database. antiSMASH v8^54^ was used to predict the BGCs in WAC276. BiG-SCAPE v2.0 cluster was used to further group the BGCs from all strains into Gene Cluster Families (GCF)(Fig S2A). Cutoff value was 0.5 and singletons were included. A presence/absence heatmap of all GCFs was generated and used to calculate the percent of shared BGCs as (Number of BGCs present in both row strain and column strain/Number of unique BGCs in row strain) (Table S2). Gene cluster comparisons of select strains were analyzed and visualized using Clinker with best links only and an identity threshold of 80%^99^.

### Planktonic Assays

*Candida albicans* ATCC90028 and *Cryptococcus neoformans* H99 were grown in 3 ml of Yeast Extract-Peptone-Dextrose (YPD) overnight at 30°C. Serial dilutions of compounds or EVs were made in 384-well plates with YP with either 2% glucose or 2% glycerol as a carbon source. Dilutions of the overnight cultures of yeast were then made in YPglucose or YPglycerol and added to the wells, final concentration 5x10^4^cells/mL, 2.5x10^3^/well. Plates were then incubated overnight at 37°C. Following incubation, OD_600_ measurements were taken of all plates (Synergy H1, Biotek). 5 µL of 0.15 µg/mL of resazurin was then added to each well and fluorescence (ex550, em590) measured after 1-3 h, once controls were visibly pink.

### Biofilm Assays

Biofilms were formed by diluting *C. albicans* and *C. neoformans* overnights in YPD to 10^7^ cfu/mL, with 100 µL of the diluted overnights seeded in each of the internal 6x10 wells of separate 96-well cell tissue treated plates. The outermost cells were filled with 100 µL of PBS and plates incubated overnight at 37°C. Treatments were prepared in 100 uL of YPD in a separate 96-well plate. Biofilms were very gently washed with 2x100 µL of PBS, and then treatments added and incubated overnight at 37°C. For the end point, biofilms were gently washed with 2x100 µL of PBS. For *C. neoformans*, 100 µL of PBS was then added and OD_600_ read for biofilm density. For *C. albicans*, 100 µL of 0.015 µg/mL of resazurin in PBS was added to each well and fluorescence (ex550, em590) was measured every 30 minutes. Inhibition was calculated from the time point where untreated controls had reached 70% of their maximum reading to ensure resazurin readings were in the linear response range.

### Hyphae Formation Microscopy

Overnight cultures of *C. albicans* ATCC90028 at 30°C were diluted to 5x10^4^cfu/mL in YPD and treated in triplicate in 96 well plates with compounds or EVs. Plates were incubated at 37°C and each well imaged in three locations every hour by Incucyte Live-Cell Analysis Instrument.

### Zones of Inhibition on Agar Plates

Overnight culture of *C. albicans* ATCC90028, SC5315, and *C. neoformans* H99 were grown in YPD at 30C. *C. albicans* cultures were diluted 1:100, *C. neorformans* 1:50 in YPglycerol and YPglycerol agar plates briefly flooded with the cell suspension. Liquid was then removed, and agar plates allowed to dry for 30 min open in a biosafety cabinet. 2 uL of serial dilutions of purified compounds or WAC276 EVs were then applied to the surface of the agar, and the agar pricked to mark spot location. Plates were incubated overnight at 30°C, and imaged at 300DPI using an Epson V850 scanner. Images were processed with Fiji ImageJ^100^, taking segment plot profiles of each zone at three different angles, measuring the diameter in pixels and defining the zone boundary as the point at which gray value was at 50% between full growth lawns and maximum inhibition. These were then converted to cm, and area of zones calculated and plotted against log of the amount of candicidin and/or antimycin applied using GraphPad Prism v11.0.2, and simple linear regressions fit.

### Treatment Loss in Biofilm Conditions Quantification Assay

*Candida albicans* ATCC 90028 cultures were grown in Yeast Extract-Peptone-Dextrose (YPD) medium at 30°C. Biofilms were seeded in 96 well flat-bottom tissue culture plates using 100 µl of 10^7^ cfu/ml overnight in YPD media and incubated for 18 hours at 37 °C. Biofilms were then gently washed with 2x 100 µl PBS before adding 100 µl of different treatments in triplicate. Treated biofilms were then incubated for an additional 3 hours. 50 uL of the treatment supernatant was then removed, and the remaining supernatant and biofilm sonicated for 5 min and vortexed for 1 min for two rounds. Disrupted biofilms were then resuspended with pipette tips and transferred to microfuge tubes, and again sonicated for 5 min and vortex for 1 min. Cells were pelleted at 5000xg for 5 min. Supernatant was removed and extracted with final 75% methanol for LC/MS-MS compound quantification.

### Biofilm Fluorescence Microscopy

During purification of a batch of WAC276 EVs, concentrated >10kDa supernatant was labeled with 10 ug/mL Alexa Fluor 555 NHS ester adding pH 8.5 sodium borate to final concentration 0.05 M and incubating for 1 h at room temperature. Labeled EVs were then purified via size exclusion chromatography and stored aliquoted at -80°C. Overnight biofilms of *C. albicans* ATCC90028 in YPD were washed and treated in duplicate with 50% WAC276 EV-555 in YPD or control cells treated with 50% PBS in YPD for 15 min, 1 h, and 3 h. After treatment biofilms were washed twice with PBS, and using a pipette tip cells from the biofilm in the centre of the well were scraped up and transferred to a microscopy slide. Cells were imaged on a Zeiss Axio Imager.M1, with Zen 2.3 (blue edition) software, for DIC and DsRed fluorescence. All images were taken with the same exposure conditions and then adjusted for brightness and contrast with identical settings. 10 fields of view were taken across the slides for each replicate.

## ACKNOWLEDGEMENTS

Funding: Canadian Institutes of Health Research, K.J.M Fellowship MFE-176478, L.E.C. Foundation grant (FDN-154288), J.R.N grant MID-406688. L.E.C. is a Canada Research Chair (Tier 1) in Microbial Genomics C Infectious Disease (CRC-2023-00184), and co-director of the CIFAR Fungal Kingdom: Threats C Opportunities program. K.J.M is supported by a career transition award from the University of Toronto, Emerging C Pandemic Infections Consortium.

## SUPPLEMENTARY FIGURES AND INFORMATION

### SUPPLEMENTARY METHODS

Planktonic assay strains: *S. cerevisiae* Y7092 (kind gift from David Andrews), *C. auris* 0387 Clinical isolate (CDC), *C. glabrata* BG2, *C. tropicalis* 1685 (Li et al 2017), *C. albicans* SN95

Li, Robbins, O’Meara, Cowen. Molecular Microbiology 2017. 103:635-656 doi 10.1111/mmi.13578 Double Knock-Out Of Candicidin And Antimycin Production

To generate an *ΔfscC* mutant for double knock-out, WAC276 genomic DNA was extracted using the DNeasy Blood C Tissue Kit (Qiagen). Two 3 kb regions corresponding to the 5’ and 3’ ends of *fscC* were amplified from genomic DNA using Phusion Polymerase (Thermo Fisher) (all primer sequences are available in Additional file X: Table X). The resulting PCR products were sequentially cloned into the HindIII site of the suicide plasmid pOJ260 (Apra^R^) to generate pOJ-fscC2. A hygromycin resistance gene (Hyg^R^) was amplified from pIJ10770 and cloned into pOJ-fscC2 at a HindIII site located between the 5’ and 3’ *fscC* flanking regions to generate pOJ-fscC-hyg. Each construct was verified by sequencing. Following successful selection of WAC276 exconjugants with hygromycin resistance, *antC* was disrupted using pOJ-antC as described in main methods.

### Biofilm Processing

Compound concentrations were considered from three compartments of each biofilm sample: Supernatant, Matrix, and Cell Pellet. Following the three-hour incubation in treated media, Supernatant samples were collected by taking 10 µl of each sample from the middle of the well careful not to disturb the biofilm. An additional 40 µl was gently removed from each sample. The remaining 50 µl of media was used to resuspend each biofilm. To disturb the biofilm, each plate was twice sonicated for 5 minutes, and gently vortexed for 1 minute, before aspirating with a pipette tip and transferring all contents to 1.5 ml microtubes. Each microtube underwent an additional 5 minute sonicated and 1 minute vortex to ensure separation of cells and matrix. Samples were then spun for 5 minutes at 5000xg to pellet cells out of the matrix. 10 µl of supernatant was then removed for Matrix LC/MS-MS analysis. The remaining supernatant was removed, and the cell pellet was washed with 1 ml of PBS. The cell pellet was then resuspended in 30 µl of 75% methanol and incubated for 1 hour to establish the Cell Pellet sample group.

**Fig S1.**
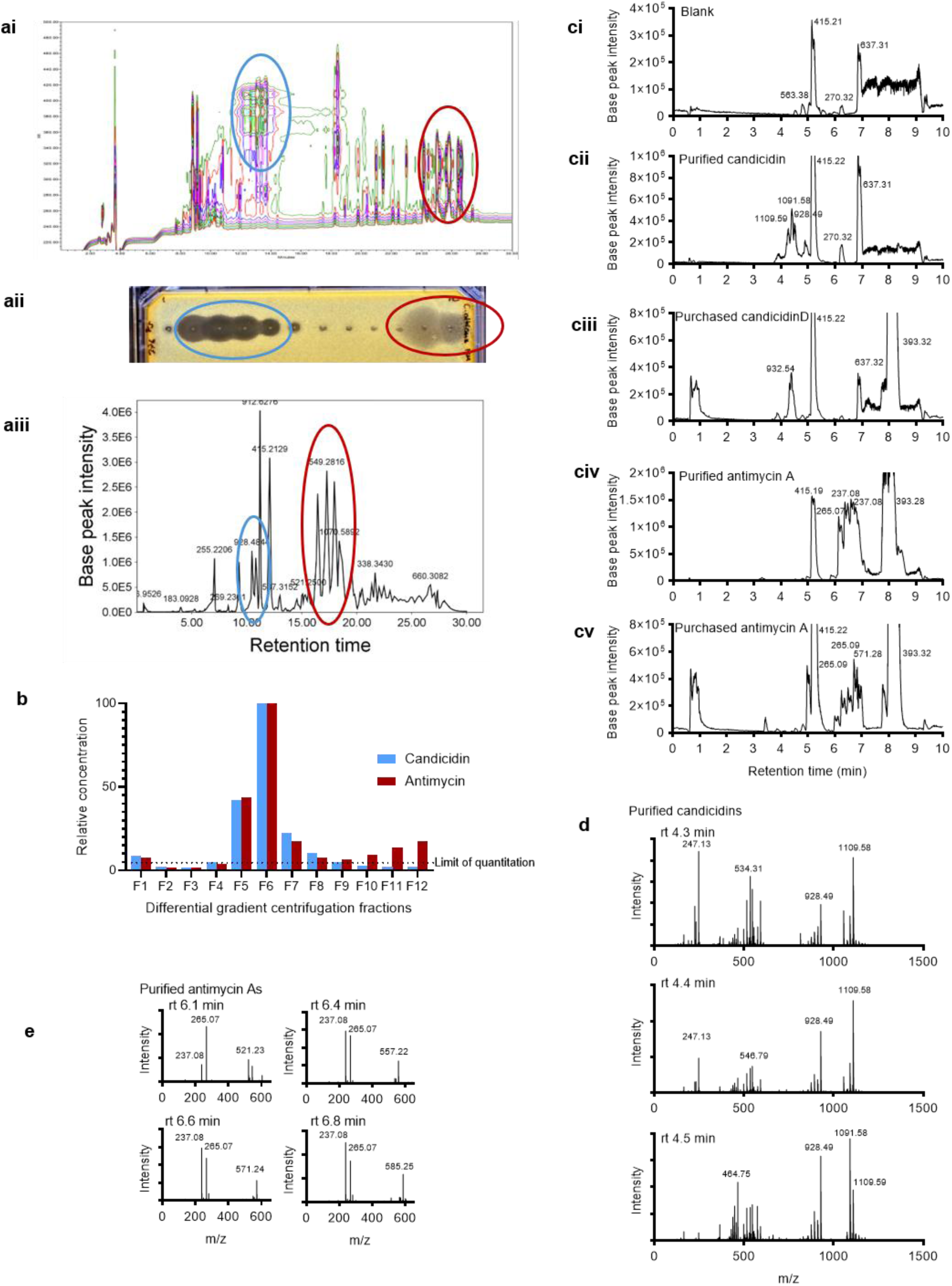
A)i) PDA surface plot of absorbance from HPLC C18 separation of WAC276 EV enriched supernatants. Blue oval highlights candicidin absorbance peaks, Red oval highlights antimycin A absorbance peaks ii) Zones of inhibition on Candida albicans lawn from fractions taken from HPLC separation, blue highlights fractions candicidin detected in, red highlights fractions antimycin As detected in, iii) base peak mass chromatogram of WAC276 EVs, ESI +ve, Q-TOF, blue highlights candicidin peaks, red highlights antimycin A peaks B) Candicidin and antimycin relative concentrations in fractions of differential gradient centrifugation layers on size exclusion chromatograph purified WAC276 EVs. Concentrations were quantified by LC-MS/MS and are relative to the F6, containing the greatest amount of candicidins and antimycins. C) Mass chromatographs for purification of candicidins and antimycins from WAC276 i) blank solvent injection, ii) purified candicidins (elute rt 4.2-4.6min), iii) purchased candicidin standard (Toku-E Company), iv) purified antimycin As (elute rt 6-7min), v) purchased antimycin A (Thermo Scientific). D) Spectrums of purified candicidin peaks (Cii) at rt 4.3, 4.4, and 4.5min. E) Spectrums of purified antimycin As (Civ) at rt 6.1, 6.4, 6.6, and 6.8 min.

**Fig S2.**
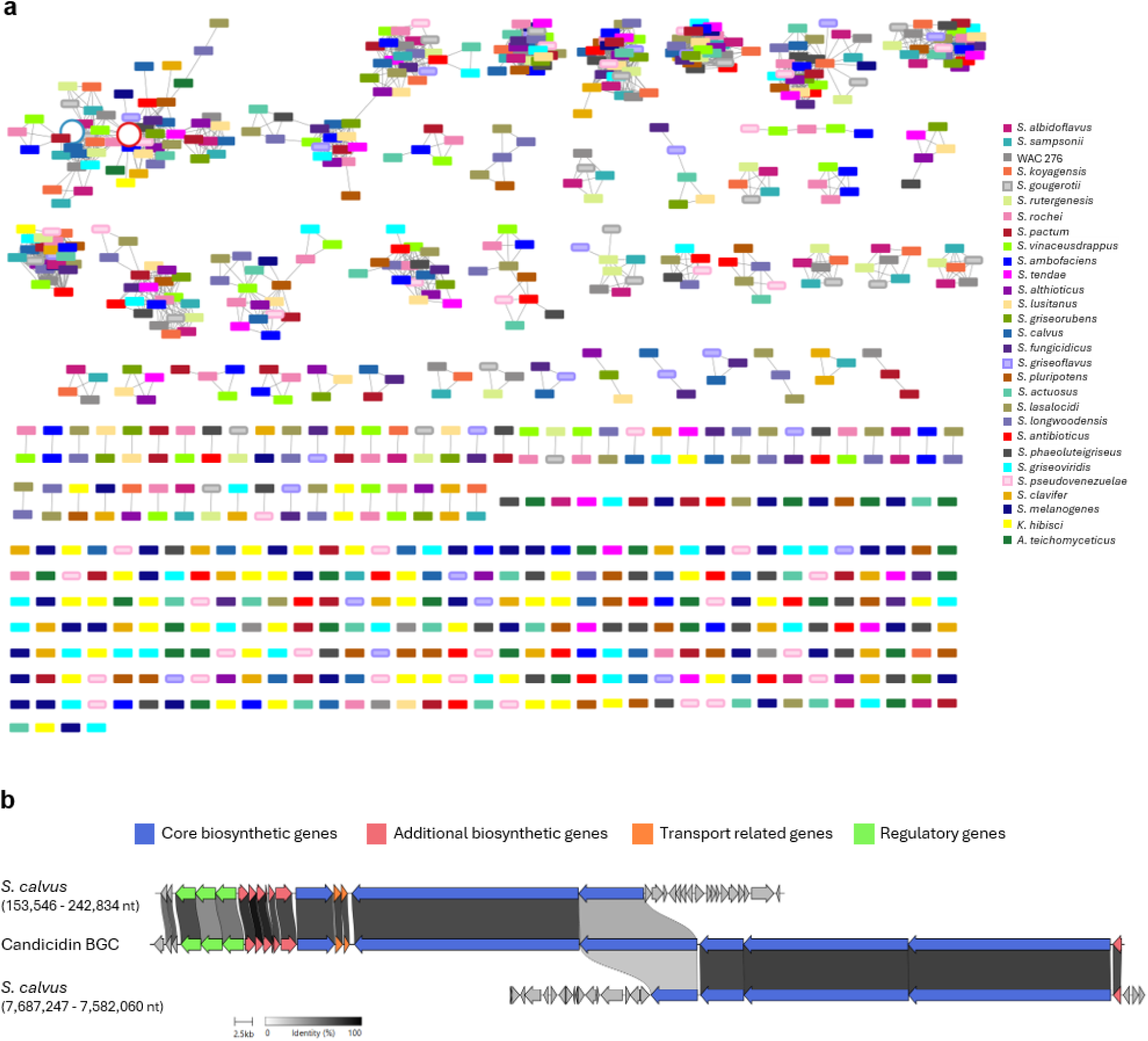
A) BiG-SCAPE network of 28 selected strains and WAC276. A cutoff of 0.5 was used. Each color represents a different strain. White circle with red outline represents antimycin BGC, white circle with blue outline represents candicidin BGC. B) Clinker alignment of the candicidin BGC with the predicted candicidin BGCs found in *S. calvus*. Identity threshold of 30% used and best links only shown. Genes colored by predicted function. Nucleotide location of each predicted BGC is included.

**Fig S3.**
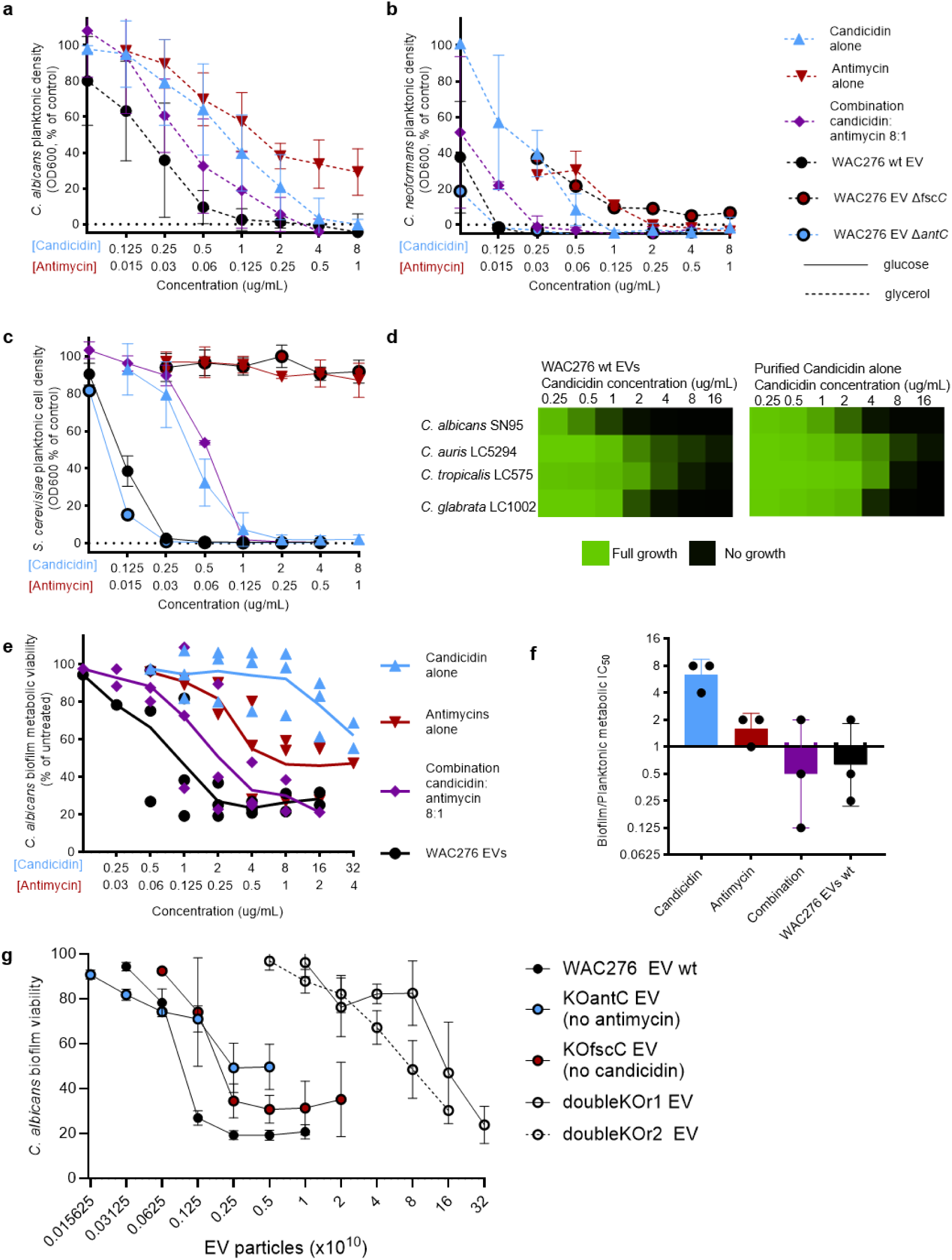
A) 37C, glycerol, B) 37C, glycerol, C) 30C, glucose, D) 30C, glucose, E) biofilm triplicate biological replicate, F) Ratio of biofilm to planktonic metabolic IC50 (concentration at which 50% of maximum effect is seen, G) Biofilm activity of EVs based on particle number, doubleKO replicate 1, doubleKO replicate 2

**Fig S4.**
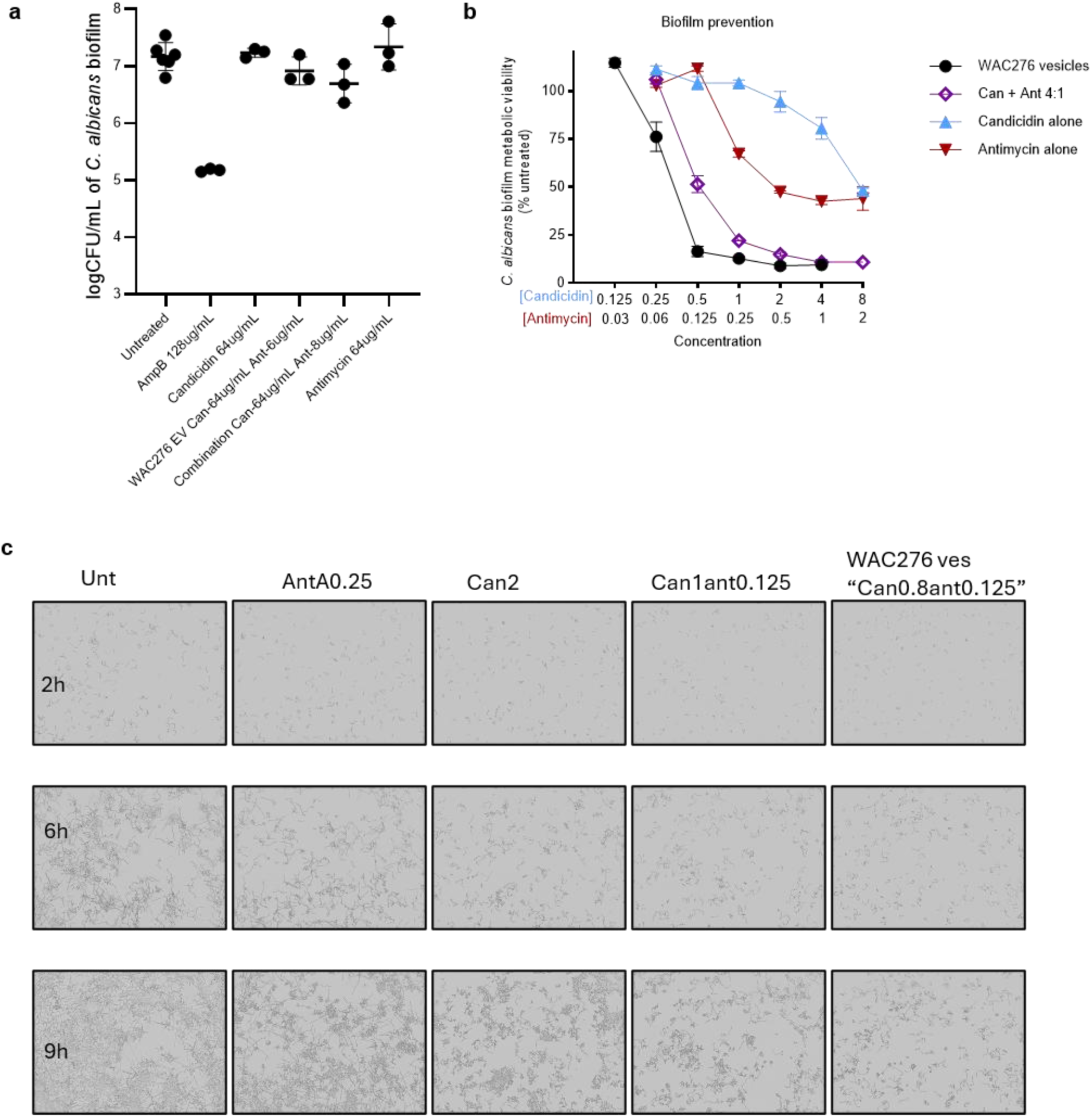
A) Killing assay of mature *C. albicans* 90028 biofilms. After 24 h treatment, biofilms were washed, sonicated, and serially diluted and plated on YPD. CFU were counted after 48 h. B) Biofilm prevention of *C. albicans* 90028 biofilms. *C. albicans* were allowed to settle in 96 well plates for 2 h, then non-adherent cells washed 2xPBS, and treatments applied in YPD for 24 h, then biofilm viability assessed with resazurin. C) Hyphae formation and cell replication at 2, 6, and 9 h of *C. albicans* 90028 in YPD at 37°C

**Fig S5.**
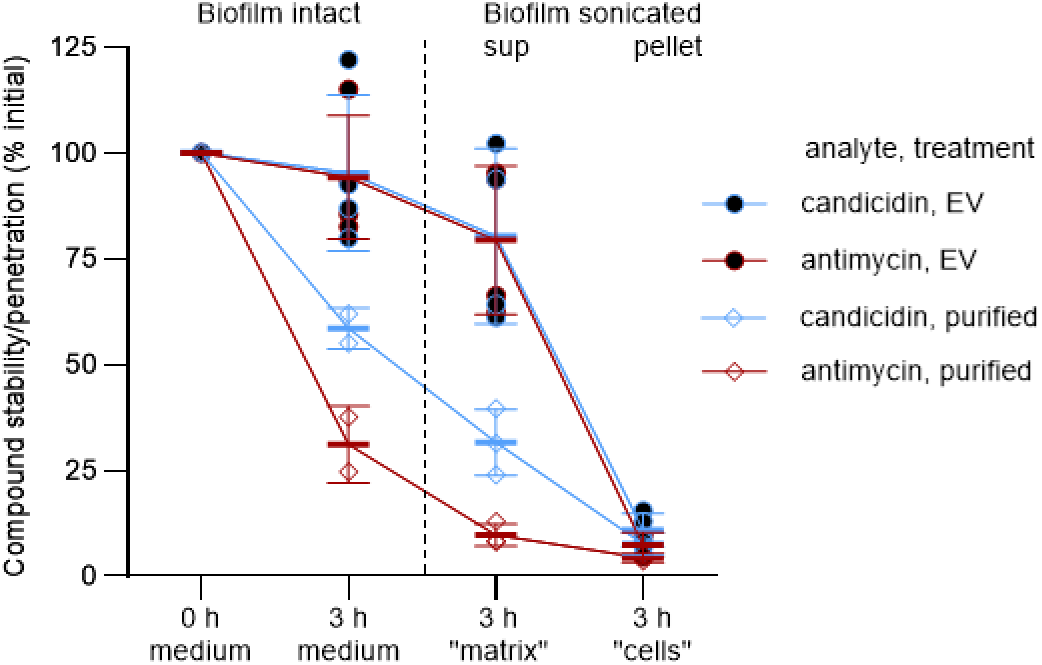
Quantification of candicidin and antimycin after WAC276 EVs or purified compounds were applied to biofilms in 96 well plates. 0 h in the YPD medium was normalized to 100%.

