## Supplementary material for "Extracellular vesicle packaging of a supercluster-encoded antimicrobial combination with enhanced potency against fungal biofilms": Table S3

| Primers for generating pOJ-fscC-hyg |  |  |
| --- | --- | --- |
| Primer name | Target | Sequence (5' - 3') |
| fscC_up_F | 5' end of fscC in WAC276 genome | TAAAACGACGGCCAGTGCCACCACCGACCTTCGCCGCACCCGG |
| fscC_up_R | 5' end of fscC in WAC276 genome | CGCGGCCGCGGATCCTCTAGAAGCTTTTGCCCATGACGACGTGCTCGCCG |
| fscC_down_F | 3' end of fscC in WAC276 genome | GGCTCGGCGAGCACGTCGTCATGGGCAAAAGCTTAAGGCGTGGAGAAGGTGACCGAGGCCCTGCGGG |
| fscC_down_R | 3' end of fscC in WAC276 genome | GGCCGCGGATCCTCTAGAAGCTAGAGTCCGAGTTCGTTGTCGAGCATG |
| hyg_F | Hygromycin resistance cassette from pIJ10770 | GAGCACGTCGTCATGGGCAAAGCCACGCGGCGGTGTCGTC |
| hyg_R | Hygromycin resistance cassette from pIJ10770 | GGTCACCTTCTCCACGCCTTAGCACGTCCCCATGCGCTCC |
| Primers for generating pOJ-antC |  |  |
| Primer name | Target | Sequence (5' - 3') |
| antC_F | 5' antC region in WAC276 genome | TAAAACGACGGCCAGTGCCAGATCTGGTCCTCGGGCTCGCC |
| antC_R | 5' antC region in WAC276 genome | CTAGAGTCGACCTGCAGCCCACCGTCACCCCCTCCTCGTCG |
| Primers to confirm ΔfscC genotype |  |  |
| Primer name | Target | Sequence (5' - 3') |
| fscC_F | Upstream left fscC homology region in WAC276 genome | ACGACTCCGACCTCGACACCAC |
| hyg_R | Hygromycin resistance cassette | GCCGAACAGCTTGATGACCGGG |
| hyg_F | Hygromycin resistance cassette | GCGACCTGCACGGGACCAACAT |
| fscC_R | Downstream right fscC homology region in WAC276 genome | GCGTCAGGTCGAGAACGGGCAG |
| Primers to confirm ΔantC genotype |  |  |
| Primer name | Target | Sequence (5' - 3') |
| antC_F | 1kb antC fragment from WAC276 genome | ACGCGGAACTGAACGCAGAGGC |
| pOJ_R | pOJ-antC backbone adjacent to 5' antC fragment | GCTTCCGGCTCGTATGTTGTGTGG |
| pOJ_F | pOJ-antC backbone adjacent to 3' antC fragment | CGCCATTCGCCATTCAGGCTGC |
| antC_R | 1kb antC fragment from WAC276 genome | GAAGGCCAGCCAGACGGTGTCC |
